# A cancer-associated polymorphism in ESCRT-III disrupts the abscission checkpoint and promotes genome instability

**DOI:** 10.1101/361659

**Authors:** Jessica B.A. Sadler, Dawn M. Wenzel, Lauren K. Williams, Marta Guindo-Martínez, Steven L. Alam, Josep Maria Mercader, David Torrents, Katharine S. Ullman, Wesley I. Sundquist, Juan Martin-Serrano

## Abstract

Cytokinetic abscission facilitates the irreversible separation of daughter cells. This process requires the Endosomal Sorting Complexes Required for Transport (ESCRT) machinery and is tightly regulated by Charged Multivesicular body Protein 4C (CHMP4C), an ESCRT-III subunit that engages the abscission checkpoint (NoCut) in response to mitotic problems such as persisting chromatin bridges within the midbody. Importantly, a human polymorphism in CHMP4C^T232^ (rs35094336), increases cancer susceptibility. Here, we explain the structural and functional basis for this cancer association: the CHMP4C^T232^ allele unwinds the C-terminal helix of CHMP4C, impairs binding to the early-acting ESCRT factor ALIX, and disrupts the abscission checkpoint. Cells expressing CHMP4C^T232^ exhibit increased levels of DNA damage and are sensitized to several conditions that increase chromosome mis-segregation, including DNA replication stress, inhibition of the mitotic checkpoint, and loss of p53. Our data demonstrate the biological importance of the abscission checkpoint, and suggest that dysregulation of abscission by CHMP4C^T232^ may synergize with oncogene-induced mitotic stress to promote genomic instability and tumorigenesis.

**Significance Statement:** The final step of cell division, abscission, is temporally regulated by the Aurora B kinase and CHMP4C in a conserved pathway called the abscission checkpoint which arrests abscission in the presence of lingering mitotic problems. Despite extensive study, the physiological importance of this pathway to human health has remained elusive. We now demonstrate that a cancer predisposing polymorphism in CHMP4C disrupts the abscission checkpoint and results in DNA damage accumulation. Moreover, deficits in this checkpoint synergize with p53 loss and generate aneuploidy under stress conditions that increase the frequency of chromosome missegregation. Therefore, cells expressing the cancer-associated polymorphism in CHMP4C are genetically unstable, thus suggesting a novel oncogenic mechanism that may involve the dysregulation of abscission.

## Introduction

Cytokinetic abscission is the final stage of cell division when the newly formed daughter cells are irreversibly separated. Abscission is a multi-step process that culminates in the resolution of the midbody, the thin intercellular bridge that connects dividing cells following mitosis (1-3). The final membrane fission step of abscission is mediated by the Endosomal Sorting Complexes Required for Transport (ESCRT) pathway (4-7). The ESCRT machinery comprises membrane-specific adaptors and five core factors/complexes (ALIX, ESCRT-I, ESCRT-II, ESCRT-III and VPS4), which are recruited sequentially (8-10). During cytokinesis, the midbody adaptor protein CEP55 initially recruits the early-acting ESCRT factors ALIX and ESCRT-I (4, 5, 11, 12). These factors, in turn, promote the recruitment and polymerization of essential ESCRT-III subunits such as CHMP4B, to form filaments within the midbody. These membrane-associated filaments collaborate with the AAA ATPase VPS4 to constrict and sever the midbody (4-6, 11, 13).

Abscission is tightly coordinated with earlier stages of mitosis to ensure faithful inheritance of genetic material during cell division (14). In particular, cytokinetic abscission is temporally regulated by a conserved mechanism known as the abscission checkpoint (NoCut in yeast), which delays abscission in response to mitotic problems such as incomplete nuclear pore reformation or chromatin bridges within the midbody (15-19). The abscission checkpoint is governed by the master regulator, Aurora B kinase, which inhibits ESCRT-III activity in response to mitotic problems. Two key intersecting signaling nodes within this pathway are the ESCRT-III subunit CHMP4C and the regulatory ULK3 kinase. CHMP4C is a specialized ESCRT-III subunit that is dispensable for cytokinetic membrane fission, viral budding and endosomal sorting, but plays an essential role in executing the abscission checkpoint (20, 21). CHMP4C is directly phosphorylated by Aurora B and is further phosphorylated by ULK3, which also phosphorylates other ESCRT-III subunits such as IST1. CHMP4C phosphorylation and ULK3 activity, together with the actions of other ESCRT-III-associated factors such as ANCHR, collectively prevent ESCRT-III polymerization and sequester VPS4 away from abscission sites, thereby delaying abscission (20, 22, 23).

Despite recent advances in identifying key components of the abscission checkpoint, the biological functions of the checkpoint and its contributions to human health are not yet known. Here, we have addressed these questions by analyzing the biochemical and abscission checkpoint activities of rs35094336, a human *CHMP4C* polymorphism (minor allele frequency [MAF]=0.04) associated with increased susceptibility to ovarian cancer (24). rs35094336 encodes an amino acid substitution of A232 (CHMP4C^A232^; reference allele) to T232 (CHMP4C^T232^; risk allele). Here, we show that the A232T substitution induces structural changes that impair ALIX binding and that cells expressing the CHMP4C^T232^ risk allele lack an abscission checkpoint and accumulate genetic damage. The CHMP4C^T232^ allele also sensitizes cells to chromosome mis-segregation and induces aneuploidy when the spindle assembly checkpoint is weakened. These observations demonstrate the importance of the abscission checkpoint in maintaining genetic stability and suggest a novel oncogenic mechanism in which disruption of the abscission checkpoint by CHMP4C^T232^ may contribute to tumorigenesis by synergizing with oncogenic mutations that increase mitotic stress.

## Results

### CHMP4C^T232^ is associated with multiple cancer types

The CHMP4C^**T232**^ allele was initially identified in a meta-analysis of two genome wide association studies (GWAS) of single nucleotide polymorphisms associated with ovarian cancer (24). To test for an association of the CHMP4C^T232^ polymorphism with other cancers, we mined data from 337,208 individuals in the UK Biobank search engine (25). Our analysis of this independent cohort confirmed the previously identified association with ovarian cancer, and revealed statistically significant associations with multiple other types of cancer, including male genital tract, prostate, and skin (Table 1). Although the odds ratios for these associations are relatively modest (1.04-1.17), association of the variant with increased risk for multiple different cancers suggests that this allele could be involved in a general pathway toward genetic instability and tumorigenesis.

**Table 1.** Significant association results for the rs35094336 genetic variant with multiple cancer types. 337,208 individuals from UKBiobank were analyzed. Results obtained from Global Biobank Engine, Stanford, CA (URL: http://gbe.stanford.edu/) [October, 2017]. The results show the association with the A (rs35094336) risk allele (CHMP4C^T232^) compared to the G (CHMP4C^A232^) reference allele.

| Cancer Type | OR (95% CI) | P-value | Cases | Cancer code |
| --- | --- | --- | --- | --- |
| Family Prostate* | 1.04 (1.01 – 1.08) | 0.007 | 34,359 | FH1044 |
| Male Genital Tract | 1.12 (1.03 – 1.23) | 0.012 | 3,449 | 1038 |
| Skin | 1.05 (1.01 – 1.09) | 0.018 | 19,170 | 1003 |
| Prostate | 1.08 (1.01 – 1.16) | 0.023 | 6,460 | 1044 |
| Non-Melanoma Skin | 1.05 (1.01 – 1.10) | 0.024 | 16,791 | 1060 |
| Ovarian | 1.17 (1.01 – 1.36) | 0.037 | 1,208 | 1039 |

\*Association with family history of prostate cancer.
OR (95% CI): Odds Ratio (95% Confidence Interval)

### CHMP4C^T232^ exhibits reduced ALIX binding

Position 232 is the penultimate CHMP4C residue, and A232 lies within a C-terminal helix that forms the ALIX binding site (26). We therefore tested whether the A232T amino acid substitution affected ALIX binding, and found that this substitution significantly reduced the interaction between full length CHMP4C and ALIX (but not CHMP4C and itself) in a yeast two-hybrid assay (Fig. 1*A*). This substitution similarly inhibited the ability of a GST-fused C-terminal CHMP4C peptide (residues 216-233) to pull down endogenous ALIX from HeLa cell lysates (Fig. S1*A*) and reduced the affinity of the terminal CHMP4C peptide for the pure recombinant ALIX Bro1 domain (residues 1-359) by 13-fold as measured in a competitive fluorescence polarization binding assay (Fig. 1*B*). In each case, we observed complete loss of ALIX binding to well-characterized control CHMP4C mutants that lacked key hydrophobic contact residues (L228A and/or W231A)(26).

**Figure. 1.**
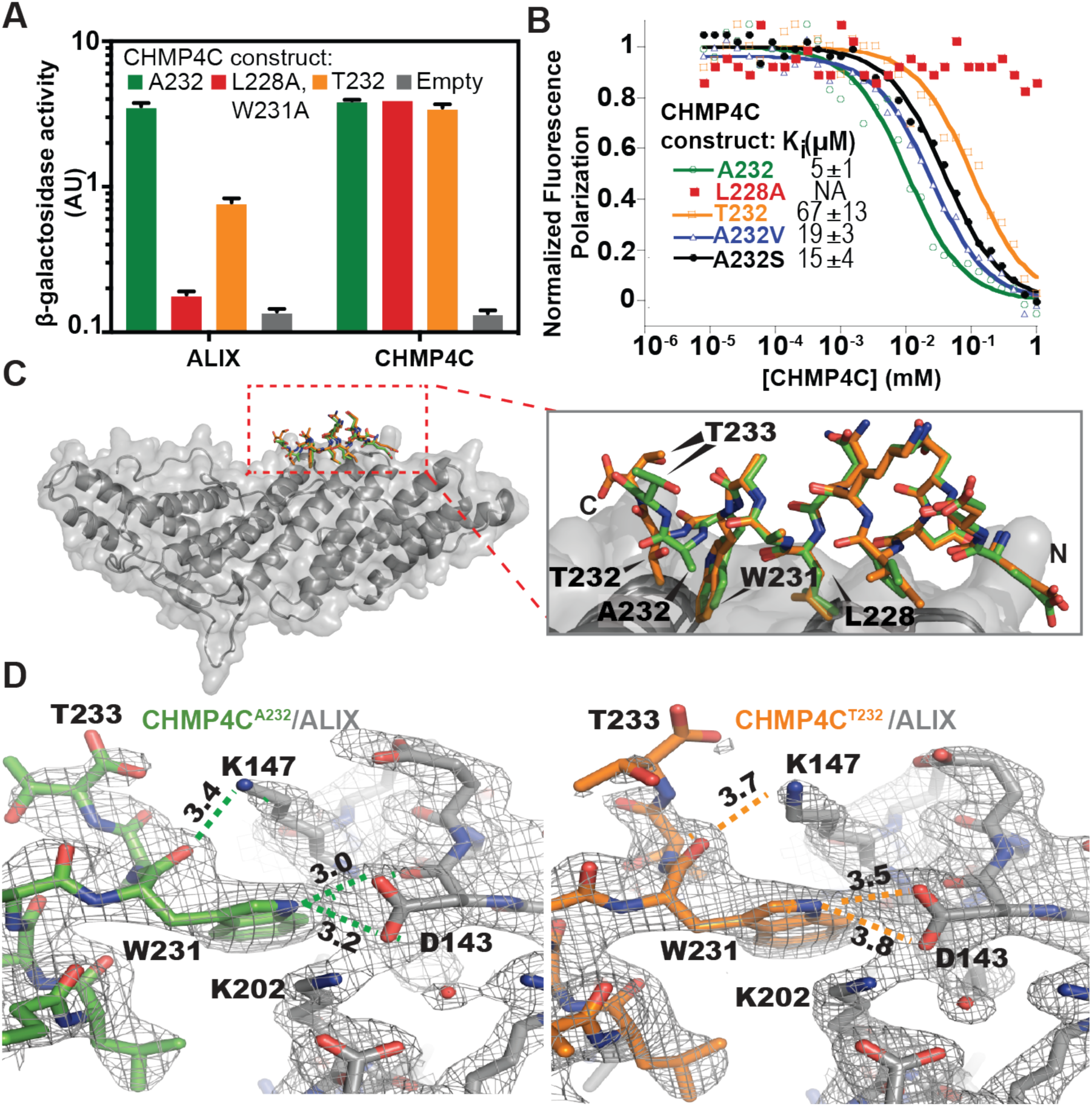
The CHMP4C A232T substitution reduces ALIX binding. **(A)** β-galactosidase activity assays of yeast co-transformed with the indicated full-length CHMP4C constructs fused to VP16 and full-length ALIX or CHMP4C fused to GAL4 (mean±SD, n=3). (**B**) Competitive fluorescence polarization binding assay with an ALIX construct spanning the Bro1 and V domains (residues 1-698) binding to fluorescently labelled CHMP4C^A232^ peptide (residues 216-233) competed with the indicated unlabeled CHMP4C peptides. Curves are from a representative experiment. K_i_ values are mean±SD. N≥7. (**C**) (Left) Superposition of ALIX Bro1 domain (grey) complexes with CHMP4C^A232^ (green) and CHMP4C^T232^ (orange) peptides. (Right) Expansion of the CHMP4C peptide superposition highlights the reduction in CHMP4C^T232^ helicity. (**D**) Binding interfaces between ALIX (grey) and CHMP4C^A232^ (left panel, green) and CHMP4C^T232^ (right panel, orange). Figures show equivalent intermolecular distances (Å) for different atoms of CHMP4C residue W231. See also Fig. S1 and Table S1.

To determine the molecular basis for this reduction in ALIX binding affinity, we determined high-resolution crystal structures of terminal CHMP4C^A232^ and CHMP4C^T232^ peptides (residues 216-233) bound to the ALIX Bro1 domain (residues 1-359) (Fig. 1*C* and Fig. S1*B*, *C* and Table S1) (26). Comparison of the structures revealed that although both peptides bound the same surface groove of the Bro1 domain, the A232T substitution disrupted several key ALIX interactions (Fig. 1*D*). Specifically, the A232T substitution unwound the C-terminal end of the terminal CHMP4C helix (Fig. 1*C*, and Fig. S1*D*), altered the position of CHMP4C residue W231, and disrupted intermolecular hydrogen bonds between the W231 carbonyl oxygen and indole nitrogen with ALIX residues D143 and K147, respectively (Fig. 1*D*). These structural analyses suggested that the A232T substitution might induce CHMP4C helix unwinding by introducing a beta-branched amino acid (which reduces helical propensity) and/or by prematurely capping the CHMP4C C-terminal helix (27). Indeed, both of these effects appeared to be operative because mutant CHMP4C peptides that selectively retained only beta-branching (CHMP4C^A232V^) or capping potential (CHMP4C^A232S^) exhibited intermediate (3-4-fold) reductions in ALIX peptide binding affinity (Fig. 1*B*). Together, these analyses demonstrate that the CHMP4C^T232^ risk allele alters the structure of the CHMP4C C-terminal helix, removes key ALIX interactions, and reduces ALIX binding affinity by more than an order of magnitude.

### ALIX-CHMP4C interactions are required for abscission checkpoint activity

ALIX is a key initiator of the cytokinetic abscission cascade (4, 5, 11, 28), and CHMP4C plays an essential role in maintaining the abscission checkpoint (4, 20, 21). We therefore tested whether abscission checkpoint activity was affected by CHMP4C mutations that impaired ALIX binding, including the CHMP4C^T232^ risk allele. In these experiments, siRNA treatment was used to deplete endogenous CHMP4C from HeLa cells engineered to stably express different siRNA-resistant HA-CHMP4C proteins (Fig. 2, and Fig. S2). Partial depletion of nuclear pore components Nup153 and Nup50 was used to activate the abscission checkpoint. As expected, control cells that expressed endogenous CHMP4C stalled during abscission, as indicated by elevated midbody connections, whereas cells depleted of CHMP4C did not exhibit elevated levels of midbody connections (20, 29) (Fig. 2*A*, Fig. S2*A*). Importantly, checkpoint activity was rescued in cells that expressed HA-CHMP4C^A232^ but not in cells that expressed HA-CHMP4C^T232^, implying that the CHMP4C^T232^ risk allele does not support the abscission checkpoint. The HA-CHMP4C^L228A,W231A^ mutant also failed to support the abscission checkpoint, further indicating that the CHMP4C-ALIX interaction is required to sustain the checkpoint. Notably, the loss of checkpoint activity for both HA-CHMP4C^L228A,W231A^ and HA-CHMP4C^T232^ was comparable to the defective response observed in cells that expressed an inactive control CHMP4C mutant lacking the amino acid insertion phosphorylated by Aurora B (HA-CHMP4C^−INS^)(20, 21). Similarly, cells expressing only HA-CHMP4C^T232^, HA-CHMP4C^L228A,W231A^ or HA-CHMP4C^−INS^ also proceeded through abscission more rapidly under normal growth conditions, implying that they were insensitive to steady-state abscission checkpoint activity, likely induced by midbody tension (23, 30) (Fig. 2*B*, Fig. S2*B*, Movies S1-S6). Despite these defects, HA-CHMP4C^A232^, HA-CHMP4C^T232^, HA-CHMP4C^L228A,W231A^ and HA-CHMP4C^−INS^ were all recruited to the midbody at normal levels, whether or not nucleoporins were depleted (Fig. 2*D*, *E*). Additionally, CHMP4C ALIX-binding mutants and wildtype CHMP4C proteins were mitotically phosphorylated at comparable levels (Fig. S2*D*), indicating that the CHMP4C mutations did not disrupt recognition by Aurora B or ULK3 kinases, and that ALIX binding is not required for these activities. These observations imply that abscission checkpoint activity requires ALIX binding to CHMP4C. We find, however, that CHMP4C midbody localization does not require ALIX binding, in contrast to a previous report (28).

**Figure. 2.**
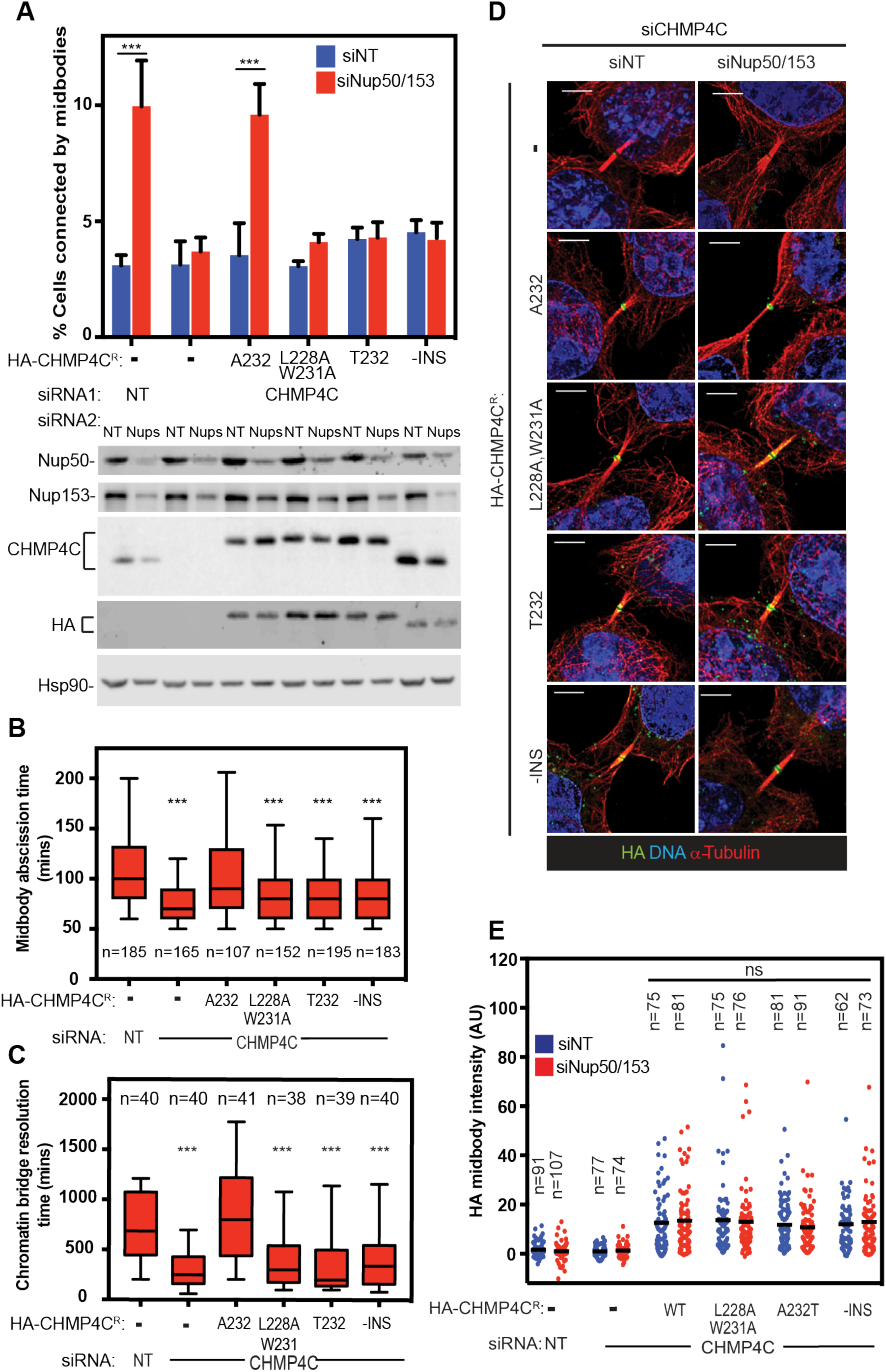
CHMP4C T232 does not support the abscission checkpoint. (**A**) HeLa cells stably expressing siRNA resistant HA-CHMP4C constructs were treated with the indicated siRNA and stained for α-tubulin. (Top) percentage of midbody-arrested cells. Data are mean±SD from >3 separate experiments with n>900 cells. (Bottom) representative immunoblots from (A). (**B**) Asynchronous HeLa mCherry-Tubulin cells stably expressing siRNA resistant HA-CHMP4C constructs were transfected with the indicated siRNA and midbody abscission times were scored from ≥3 separate experiments. Here and throughout, box edges mark the 25^th^ and 75^th^ quartiles and whiskers mark 5-95% percentile. Bars denote the median. Mean times ±SD were non-targeting (NT): 110±47 min, siCHMP4C: 75±26 min HA-CHMP4C^A232^ + siCHMP4C, 108±47 min, HA-CHMP4C^L228A,W231A^ + siCHMP4C, 86±33 min, HA-CHMP4C^T232^ + siCHMP4C, 81 ±27 min, HA-CHMP4C^−INS^ + siCHMP4C, 87±44 min. See also Movies S1-S6. (**C**) HeLa cells stably expressing siRNA resistant HA-CHMP4C constructs and YFP-Lap2β were transfected with the indicated siRNA. Resolution time of Lap2β bridges were quantified from >3 separate experiments. Mean times ±SD were nontargeting (NT): 721 ±333 min, siCHMP4C: 296±187 min HA-CHMP4C^A232^ + siCHMP4C, 853±474 min, HA-CHMP4C^L228A,W231A^ + siCHMP4C, 415±317 min, HA-CHMP4C^T232^ + siCHMP4C, 396±394 min HA-CHMP4C^−INS^ + siCHMP4C, 421 ±341 min. See also Movies S7-S12. (**D, E**) HA-CHMP4C recruitment to midbodies determined by staining for DAPI (blue), a-tubulin (red) and HA (green) from >3 separate experiments. Scale bar, 5 μm. Data in (**E**) represent the staining intensity of HA normalized to background measurements. Mean value is marked. P-values were calculated using 2-way ANOVA and Sidak’s multiple comparisons test, (**A**, **E**) or one-way ANOVA vs. control (HeLa siNT) (**B, C**), ∗∗∗=p<0.001. Western blots for B, C, are shown in Fig. S2.

To test CHMP4C^T232^ activity when the abscission checkpoint was activated by a different trigger, we used live cell imaging to measure the resolution times of intercellular chromatin bridges, as visualized using the nuclear envelope marker lamina-associated polypeptide 2β fused to YFP (YFP-LAP2β). As expected (20), chromatin bridges were resolved prematurely in CHMP4C-depleted cells compared to cells that expressed endogenous CHMP4C^A232^ (median resolution time = 250 vs 685 min, Fig. 2*C*, Fig. S2*C*, Movies S7-S12). Importantly, normal midbody resolution times were restored by expression of siRNA-resistant HA-CHMP4C^A232^ (800 min), but not by HA-CHMP4C^T232^ (200 min), HA-CHMP4C^L228A,W231A^ (300 min) or HA-CHMP4C^−INS^ (340 min). Thus, cells expressing the cancer-associated CHMP4C^T232^ risk allele lack an appropriate abscission checkpoint response under multiple different conditions that activate this checkpoint.

### Disruption of the abscission checkpoint leads to accumulation of DNA damage

Although the biological consequences of abscission checkpoint loss are not well understood, increased DNA damage is one possible outcome (20). We therefore compared DNA damage accumulation in cells that expressed the different CHMP4C mutants. In these experiments, CRISPR-Cas9 was used to delete the CHMP4C locus from HCT116 cells (Fig. S3), a near diploid cell line that exhibits low levels of chromosomal instability and DNA damage (31). Genetic damage was then assessed by scoring the number of nuclear foci formed by the DNA damage response marker 53BP1. As expected, cells with low levels of DNA damage predominated in the wild type cultures (HCT116^wt^) (<2 foci/cell) (Fig. 3*A*, *B*). In contrast, cells lacking CHMP4C (HCT116^δCHMP4C^) exhibited heightened DNA damage (>6 foci/cell) with significantly greater frequency. Crucially, DNA damage in the HCT116^δCHMP4C^ cells was reduced to control levels upon re-expression of HA-CHMP4C^A232^, but not HA-CHMP4C^T232^, HA-CHMP4C^L228A,W231A^ or HA-CHMP4C^−INS^. Hence, loss of the CHMP4C-dependent abscission checkpoint increases the accumulation of 53BP1-associated DNA damage foci.

**Figure. 3.**
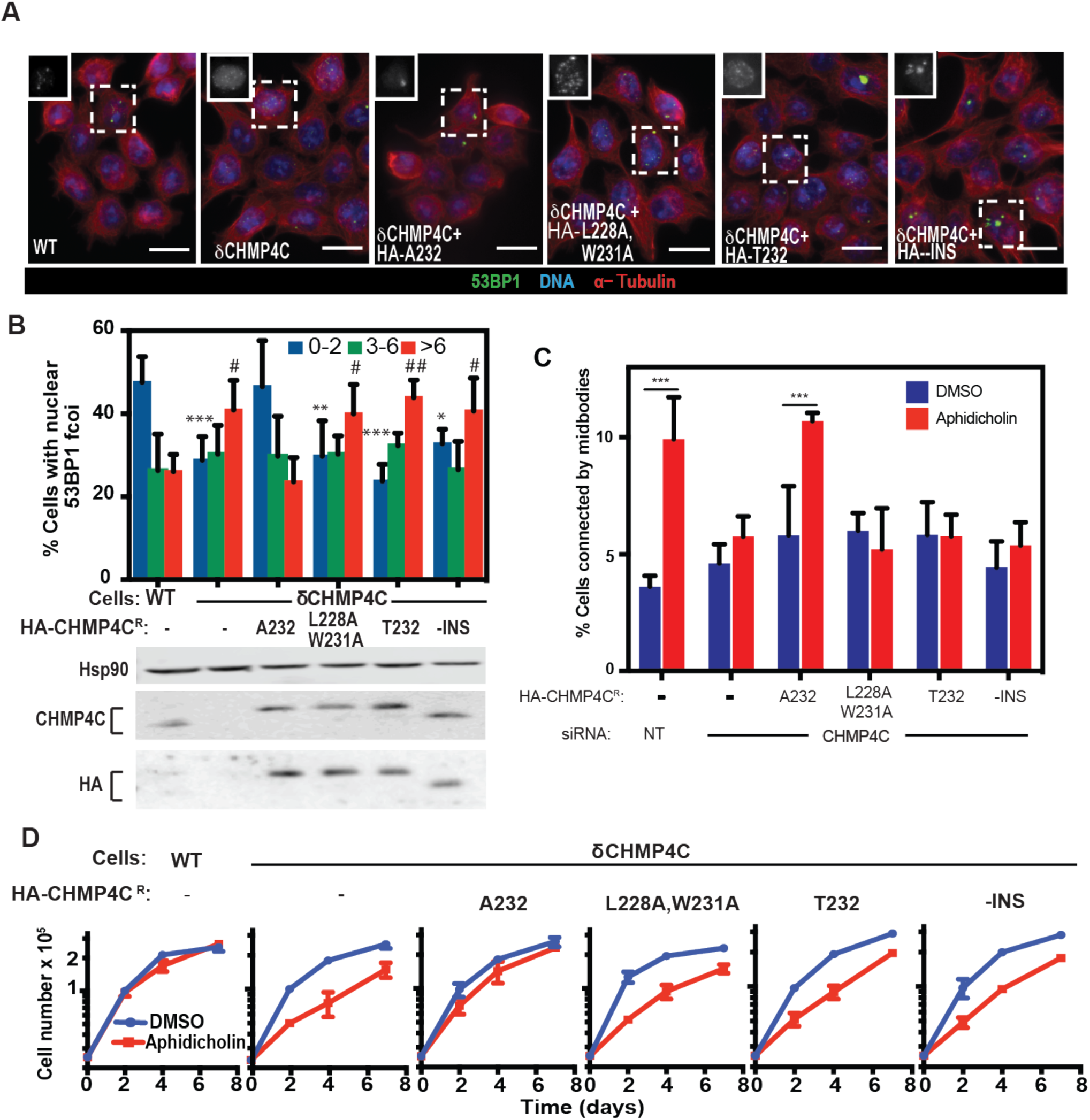
Cells lacking the abscission checkpoint exhibit elevated genome damage and are sensitized to replication stress. (**A**) HCT116 cells with (WT) or without endogenous CHMP4C (δHMP4C) expressing the indicated HA-CHMP4C construct were stained for 53BP1 (green), α-tubulin (red) and DAPI (blue). Scale bars, 20 μm. Insets show grey scale images of boxed cells. (**B**) Numbers of 53BP1 foci/cell were determined from 3 independent experiments and binned into the designated categories. Shown are mean±SD from >900 cells. P-value calculated using 2-way ANOVA and Sidak’s multiple comparisons test comparing each category to control (WT), ∗= 0-2 foci, #= >6 foci, ∗/#P<0.05 ∗∗/##=P<0.005, ∗∗∗/###=P<0.001. (**C**) Cells stably expressing the indicated HA-CHMP4C construct were treated with the indicated siRNA for 48 hours, then with DMSO or 30 nM aphidicolin for 24 hours. The number of cells connected by midbodies was scored. (**D**) HCT116^WT^ or HCT116δ^CHMP4C^ cells expressing the indicated HA-CHMP4C constructs were cultured in the continuous presence of DMSO (blue) or 30 nM aphidicolin (red) and cell numbers were determined at the indicated time points. Plotted are mean±SD from 3 independent experiments. See also Fig. S7*A.*

### CHMP4C is not required for maintenance of nuclear integrity, DNA damage responses nor mitotic checkpoint signaling

To determine whether observed phenotypes were specifically due to abscission checkpoint failure, we examined other mechanisms that might underlie the elevated levels of DNA damage in HCT116^δCHMP4C^ cells. We ruled out ESCRT-dependent loss of nuclear envelope integrity (32, 33) because nuclear envelope compartmentalization was not compromised during telophase in cells lacking CHMP4C, as assayed by nuclear morphology and retention of a GFP-NLS reporter (Fig. S4 and Movies S13-S15). Similarly, loss of CHMP4C did not globally impair DNA damage responses because the efficiency of G2/M cell cycle arrest in response to genotoxic stress induced by the DNA cross-linker mitomycin C was normal in HCT116^δCHMP4C^ cells (Fig. S5). Furthermore, in contrast to another report (34), we did not observe a failure of HCT116^δCHMP4C^ cells to arrest in response to spindle poisons such as nocodazole, indicating that the spindle assembly checkpoint remains largely intact in these cells (Fig. S6). Therefore, the increased DNA damage in cells lacking CHMP4C activity and a functional abscission checkpoint does not reflect loss of nuclear integrity, improper DNA damage responses or defective mitotic spindle assembly checkpoint signaling but rather a loss of the abscission checkpoint.

### CHMP4C^T232^ sensitizes cells to replication stress

We next examined the possibility that cells lacking CHMP4C activities had increased levels of DNA damage because they were unable to respond properly to DNA replication stress. This is an attractive model because: 1) a significant fraction of 53BP1 nuclear bodies originate from lesions generated by DNA replication stress (35), 2) elevated replication stress triggers the abscission checkpoint in a CHMP4C-dependent manner (18) (Fig. 3*C*), and 3) the abscission checkpoint plays a key role in protecting anaphase bridges that arise from replication stress (36), thereby reducing damage when they persist during cytokinetic abscission (37). In agreement with this model, inducing replication stress with ultra-low doses (30 nM) of the DNA polymerase inhibitor aphidicolin reduced the proliferation of HCT116^δCHMP4C^ cells nearly 2-fold as compared to HCT116^WT^ cells (Fig. 3*D*). Importantly, the HCT116^δCHMP4C^ growth defect was again rescued by expression of HA-CHMP4C^A232^, but not by the abscission checkpoint defective, HA-CHMP4C^T232^, HA-CHMP4C^L228A,W231A^ or HA-CHMP4C^−INS^ mutants. Thus, the abscission checkpoint can play a protective role in cell survival when cells are subjected to increased replication stress.

### CHMP4C^T232^ sensitizes cells to chromosome mis-segregation and induces aneuploidy

To examine the functions of the abscission checkpoint in the context of another mitotic stress, we tested whether a defective abscission checkpoint also sensitized cells to weakening of the spindle assembly checkpoint (SAC), a condition that induces anaphase chromosomal segregation errors. The SAC was selectively weakened by treatment with low doses (0.1 μM) of the MPS1 kinase inhibitor reversine (38), which doubled the frequency of anaphase chromosome mis-segregation (Fig. 4*A*, Fig. S7*B*, Movies S16-19) and activated the abscission checkpoint in a CHMP4C-dependent fashion (Fig. S7*A*). Karyotyping of metaphase spreads revealed that HCT116^WT^ cells only rarely displayed extreme aneuploidy (1% of DMSO-treated cells had <37 or >48 chromosomes, Fig. 4*B*, *C*). Reversine treatment alone increased this percentage, but cells with extreme aneuploidy were still rare (7%). In contrast, DMSO treated HCT116^δCHMP4C^ cells exhibited a higher basal level of extreme aneuploidy (4%), and this percentage increased notably upon reversine treatment (23%). Re-expression of HA-CHMP4C^A232^, but not HA-CHMP4C^T232^ protected against reversine induced increases in aneuploidy (Fig. S8*A*). Importantly, reversine treatment in cells with a defective abscission checkpoint did not induce a multinucleation phenotype, but these cells did display a higher proportion of micronuclei (Figure S8*B*).

**Figure. 4.**
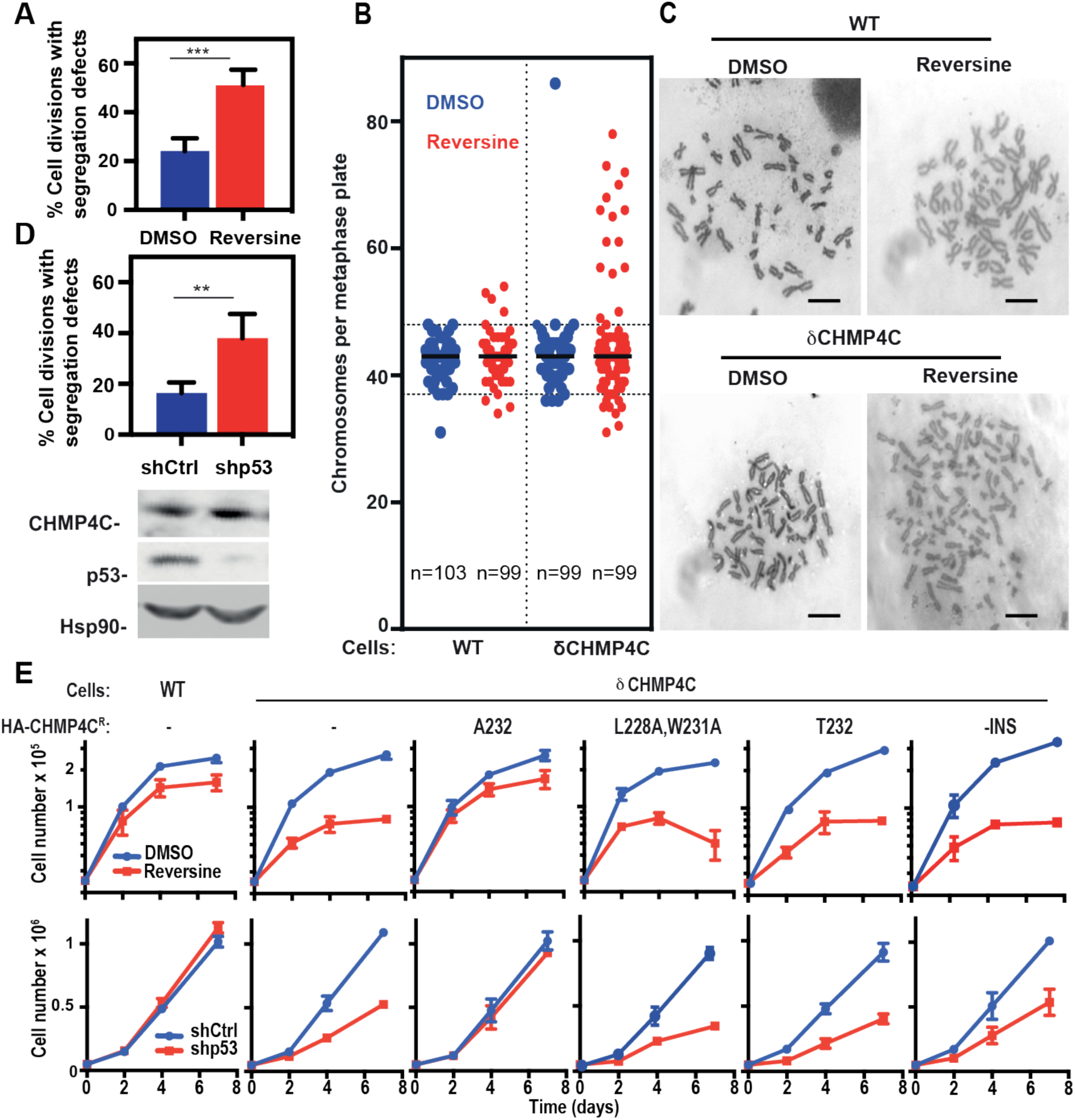
Defects in chromosome segregation and the abscission checkpoint synergize to impair cell growth. (**A**) HCT116 cells expressing histone H2B-mCherry were grown in the continuous presence of DMSO (blue) or 0.1 μM reversine (red) and anaphase segregation defects were scored. Data shown are mean±SD from n>130 cells from 3 separate experiments. See also Fig. S7*B, C*, and Movies S16-S19. (**B**) HCT116^WT^ or HCT116δ^CHMP4C^ were cultured for 48 hours in the continuous presence of DMSO (blue) or 0.1 μM reversine (red). Metaphases were enriched by overnight treatment with nocodazole, and chromosome number was determined. Plots show all data with medians marked. Extreme aneuploidy is defined as chromosome numbers above 48 or below 37 (dashed lines). Representative metaphase spreads can be seen in (**C**), scale bars, 10 μm. Data were collected from >4 independent experiments. (**D**) HCT116 cells expressing histone H2B-mCherry were depleted of p53 by shRNA treatment (shp53) and anaphase segregation defects were scored. Data are mean±SD, n>200, from 4 separate experiments across two shRNA transductions. See also Fig. S7*D*, S9*B*, *C*, and Movies S20-S23. (**E**) HCT116^WT^ or HCT116δ^CHMP4C^ cells expressing the indicated CHMP4C construct were (top panel) grown in the continuous presence of either DMSO (blue lines) or 0.1 μM reversine (red line) or (bottom panel) depleted of p53 by shRNA treatment (shCtrl, blue, shp53, red) and cell numbers were determined at the indicated time points. Data show are mean±SD from >3 independent experiments. *P*-values calculated using two-tailed unpaired Student’s f-test, ∗∗P<0.005,∗∗∗P<0.001. See also Fig. S9*A*

In a complementary set of experiments, we monitored cell growth rates in reversine-treated cultures, where the high levels of chromosomal instability and aneuploidy are detrimental to cell survival and growth (39) (Fig. 4*E*, top panel). In agreement with the karyotype analyses, reversine treatment reduced HCT116^WT^ cell growth only modestly, but reduced HCT116^δCHMP4C^ cell growth by at least 70% over a seven-day period. Robust growth in the presence of reversine was restored by expression of HA-CHMP4C^A232^, but not HA-CHMP4C^T232^, HA-CHMP4C^L228A,W231A^ or HA-CHMP4C^−INS^ (Fig. 4*E*, top panel). Thus, the CHMP4C-dependent abscission checkpoint becomes more critical for cell growth under conditions that increase chromosome segregation defects.

### CHMP4C^T232^ synergizes with p53 loss

Our observations that CHMP4C mutations can abolish the abscission checkpoint and that these mutations synergize with increases in chromosome mis-segregation raised the intriguing possibility that such mutations might also synergize with genetic alterations known to be associated with ovarian cancer development. We focused our studies on TP53, the most frequently mutated gene in a wide range of cancers, including >96% of high-grade serous ovarian tumors (40). In addition to its classical role in inducing cell cycle arrest in response to DNA damage, p53 also functions directly in DNA damage repair and chromosomal stability, and its absence is associated with increased replicative stress (41). As expected, stable depletion of p53 increased DNA damage levels as measured by increased numbers of 53BP1 foci (Fig. S9*B*, *C*) and increased frequencies of chromosomal segregation defects during anaphase (Fig. 4*D*, S7*D*, Movies S20-S23). As with reversine treatment, these defects significantly compromised HCT116 cell growth only when CHMP4C was absent (Fig. 4*E*, bottom panel, Fig. S9*A*). This synergistic effect was again reversed upon re-expression of CHMP4C^A232^ but not HA-CHMP4C^T232^, HA-CHMP4C^L228A,W231A^ or HA-CHMP4C^−INS^ (Fig. 4*E*, bottom panel, Fig. S9*A*). Thus, loss of the CHMP4C-dependent abscission checkpoint also synergizes with loss of functional p53. This effect can again be explained by an inability of cells to cope with the increased burden of chromosomal segregation defects, perhaps compounded further by dysfunction of the p53-mediated G1 checkpoint (42).

## Discussion

Our study demonstrates that the abscission checkpoint plays a critical role in human health by protecting the genome against DNA damage and chromosomal instability. We have shown that a human polymorphism in *CHMP4C,* previously associated with increased susceptibility to ovarian cancer, is also associated with increased risk for several other cancers, thus suggesting that the CHMP4C^T232^ allele contributes to tumor development in a global fashion. Importantly, cells that express the cancer-associated CHMP4C^T232^ allele show elevated levels of 53BP1 foci, suggesting that increased DNA damage may account, at least in part, for increased cancer susceptibility in individuals who carry this allele. Furthermore, cells that express CHMP4C^T232^ are particularly sensitized to genomic instability under conditions that increase the burden of chromosomal segregation defects, such as DNA replication stress and weakening of the spindle assembly checkpoint.

At the mechanistic level, the A232T substitution unwinds the C-terminal CHMP4C helix, impairs ALIX binding affinity, and leads to loss of abscission checkpoint activity. Hence, in addition to the previously documented requirements for CHMP4C phosphorylation by Aurora B and ULK3 (20, 21, 23), CHMP4C must also be able to interact with ALIX (and possibly other Bro-domain containing proteins) to support the abscission checkpoint. We found, however, that altered CHMP4C midbody localization could not explain the loss of checkpoint activity because point mutations that specifically abolished ALIX binding did not reduce CHMP4C midbody localization or mitotic phosphorylation. In contrast, others have observed reduced midbody localization of a C-terminally truncated CHMP4C construct (28). It is therefore possible that the C-terminal region of CHMP4C dictates midbody localization independently of ALIX binding, perhaps through interactions with MKLP1 or the Chromosomal Passenger Complex (20, 43). Alternatively, removal of 18 terminal CHMP4C residues could have relieved CHMP4C autoinhibition, thereby impairing midbody localization indirectly. This idea is consistent with the observation that autoinihibition of Snf7p, the yeast orthologue of CHMP4, can be relieved by removing its terminal Bro1p (ALIX) binding helix (44-46). We do not yet know for certain why CHMP4C-ALIX binding is required to support the abscission checkpoint, but one intriguing possibility is that CHMP4C binding may competitively inhibit CHMP4B from occupying its overlapping binding site on ALIX (26), thereby sustaining the abscission checkpoint by preventing nucleation of CHMP4B-containing ESCRT-III filaments within the midbody.

Another striking finding of our study is that increasing chromosomal segregation defects in cells lacking a functional abscission checkpoint specifically induces high levels of aneuploidy and chromosomal instability. Although complete inhibition of Aurora B leads to cleavage furrow regression and binucleation when chromatin is present in the intracellular bridge (17), this mechanism does not appear to explain the increases in aneuploidy when CHMP4C is impaired. Depletion of CHMP4C (or other abscission checkpoint components downstream of Aurora B such as ULK3) induces premature resolution of chromatin bridges, not furrow regression (20, 23). Moreover, reversine treatment in cells lacking CHMP4C does not lead to multinuleation, but does induce micronuclei formation. Our data are consistent with chromosomal instability resulting from chromosome breakage and refusion events, and/or the failure to re-incorporate lagging chromosomes into the main nuclei. It has been suggested that even very mild aneuploidy has consequences beyond chromosome gains or losses, resulting in DNA damage and replication stress which have severe effects on subsequent mitoses, and that even gain of a single chromosome can result in further chromosomal aberrations and complex karyotypes in subsequent cell cycles (47, 48). We suggest that DNA damage acquired over a number of cell cycles in cells lacking a functional abscission checkpoint may have cumulative effects that are ultimately detrimental in subsequent cell cycles. Consistent with this idea, damage acquired during mitosis can lead to p53-dependent quiescence in daughter cells (47, 48). Thus, the coordinated action of the abscission checkpoint and p53 may protect against aneuploidy. In this model, the abscission checkpoint provides additional time to retrieve, repair and/or protect lagging chromosomes, thereby protecting cells and preventing catastrophic DNA damage during mitosis. Disruption of the abscission checkpoint could also induce aneuploidy via a mechanism that is reminiscent of the checkpoint adaptation phenomenon, in which cells continue to proceed through the cell cycle despite not having completely resolved DNA damage arising the previous mitosis (51). Such checkpoint-adapted cells are characterized by severe chromosomal segregation defects that give rise to micronuclei containing lagging chromosomes (or chromosome fragments), which then contribute to further chromosome damage and instability in subsequent divisions.

Chromosomal segregation errors are a hallmark of many cancers and often arise in response to oncogenic mutations that increase mitotic stress (39). In particular, loss of p53 is the most common genetic abnormality in many tumor types, and can lead to increased DNA damage, chromosomal instability, and increased replicative stress (40, 41, 50). These phenotypes suggest a potential mechanism by which loss of the abscission checkpoint could contribute to genetic instability and cancer development, particularly as we observed synthetic lethality between loss of p53 and the CHMP4C^T232^ risk allele. Our data suggest the possibility that homozygous germline expression of the CHMP4C^T232^ allele - or loss of heterozygosity of CHMP4C^A232^ encoding allele expression in somatic cells where the T232 encoding allele is present - could contribute to tumorigenesis by increasing genomic instability and aneuploidy, particularly when chromosome mis-segregation events are elevated. We speculate that although an impaired abscission checkpoint combined with genetic alterations such as p53 loss is detrimental to overall cell growth, the subset of cells that ultimately survive may accumulate further adaptations that promote tumorigenicity. These cells may nevertheless remain sensitive to further perturbations of chromosome segregation or DNA damaging agents and this sensitivity could, in principle, be exploited therapeutically. In this regard, it is noteworthy that CHMP4C depletion increases the effectiveness of irradiation-induced apoptosis in human lung cancer cells (51) and that many common chemotherapeutic drugs such as Paclitaxel act, at least in part, by increasing chromosome mis-segregation (39). Hence, we speculate that such chemotherapeutics may be particularly effective in patients who carry the CHMP4C^T232^ allele.

## Methods

### Plasmids and antibodies

Details of plasmids and antibodies used in this study are described in Table S2.

### Fluorescence polarization (FP) binding experiments

Fluorescence polarization was measured using a Biotek Synergy Neo Multi-Mode plate reader (Biotek) with excitation at 485 nm and detection at 528 nm. For competitive binding experiments, the wild type CHMP4C peptide (residues 216-233) was synthesized with a non-native cysteine at the N-terminus (CQRAEEEDDDIKQLAAWAT), and labeled with Oregon Green 488 (Life Technologies/Molecular Probes 06,034) following manufacturer’s instructions. The labeled peptide was quantitated by the absorbance of Oregon Green 488 at 491 nm (extinction coefficient 83,000 cm^−1^ M^−1^ in 50 mM potassium phosphate, pH 9). Different concentrations (as determined by absorbance at 280 nm) of unlabeled, N-terminally acetylated CHMP4C peptides were titrated against a CHMP4C^A232^-ALIX Bro1-V complex created by mixing 5 μM ALIX Bro1-V and 0.5 nM fluorescently labeled CHMP4C^A232^ peptide in binding buffer (20 mM sodium phosphate pH 7.2, 150 mM NaCl, 5 mM BME, 0.01% Tween-20, and 0.2 mg/mL Bovine Serum Albumin). IC(50)s were calculated from binding curves using Kaleidagraph (Synergy Software) and converted to K(i) (54). Competitive binding curves were measured independently ≥7 times for each peptide and mean±SD K(i) values are reported. All peptides were synthesized by the University of Utah Peptide Synthesis Core Facility, and verified by mass spectrometry.

### GST Pull-Downs

GST-fused CHMP4C peptides spanning the ALIX binding helix (residues 216-233) from CHMP4C^A232^ or the CHMP4C^L228A,W231A^ or CHMP4C^T232^ mutants were purified, immobilized on glutathione-Sepharose agarose beads, and incubated with clarified HeLa cell lysates. Bound material was analysed by SDS-PAGE followed by immunoblotting or Coomassie staining. A detailed description is provided by SI Materials and Methods.

### Cell culture

HEK293T, HeLa and HCT116 cells were cultured and maintained in DMEM medium supplemented with 10% fetal bovine serum and 20 μg/mL gentamycin. To generate stable cell lines, 293T cells were transfected with retroviral packaging vectors (Table S2), MLV-GagPol/pHIV 8.1 and pHIT VSVg at a ratio of 3:2:1 for 48 hours. 293T supernatant was filtered through a 0.2 μm filter and used to transduce the indicated cell lines, with antibiotic selection carried out 48 hours later. MycoAlert (Lonza) was used to screen for mycoplasma contamination. HCT116 CRISPR cell lines were generated by transfection with retroviral Cas9 expression plasmids containing specific guide RNAs targeting CHMP4C, and full details are available in SI Materials and Methods.

### siRNA transfections

Cells were transfected with siRNA for 72 hours using Dharmafect 1 (Dharmacon) or Lipofectamine RNAiMax (Thermofisher) according to manufacturers’ instructions. Cells received two transfections, once at 0 hours and again at 48 hours. For HeLa cells, CHMP4C and non-targeting siRNA were used at 100 nM and Nup50 and Nup153 siRNA were used at 10 nM. For HCT116 cells, all siRNA was used at 10 nM. Cells were fixed or imaged 24 hours after the second siRNA transfection. For GFP-NLS nuclear fluorescence recovery experiments HCT116 cells were imaged 8 hours after the second transfection. siRNA sequences used in this study have been described (16-19) and are available in Table S2.

### Immunoblotting

Cell lysates were denatured in Laemmli buffer, resolved by SDS-PAGE, and transferred to Nitrocellulose membranes. Membranes were blocked with 5% skim milk in 0.1% Tween 20/TBS and incubated with primary antibodies in either 1% or 5% skim milk in blocking solution for 3 hours at room temperature or overnight at 4°C. Membranes were washed in 0.1% Tween 20/TBS and incubated with the corresponding secondary antibodies conjugated with HRP or near-Infrared fluorescent dyes in blocking solution for 1 hour at room temperature and washed again. Proteins were detected and quantified using a Li-Cor Odyssey Infrared scanner and software (Li-Cor Biosciences) or Image Quant LAS 400 (GE Healthcare). Details on antibodies and dilutions can be found in Table S2.

### Immunofluorescence

Cells were grown on coverslips, washed once in PBS and fixed for 10-20 minutes in ice cold methanol. Cells were blocked with 3% FCS, 0.1% Triton X-100 in PBS for 20 minutes. Primary antibodies were applied for at least 1 hour. After washing 4X with PBS, secondary antibodies were applied for 1 hour and nuclei were stained with either Hoechst or 4′,6-diamidino-2-phenylindole (DAPI). Coverslips were mounted with ProLong Gold Antifade Reagent (Invitrogen) on a microscope slide. Images were acquired using a Leica SP8 Confocal (Fig. 2*D*) or Nikon Ti-Eclipse wide-field inverted microscope (Fig. 3, S4, S2). Scoring was conducted blind. Where indicated, deconvolution was performed using HyVolution Pro-Automatic deconvolution software. Quantification of fluorescence staining intensity was carried out with ImageJ. The freehand selection tool was used to outline the region of interest and staining intensity within this area was measured. Signals were background corrected using measurements from adjacent regions. Details on antibodies and dilutions can be found in Table S2.

### Live cell imaging

Cells were seeded on glass-bottomed 24-well plates (MatTek) and transfected with siRNA or shRNA or subjected to the specified drug treatments. Imaging was carried out for 24-72 hours on a Nikon Ti-Eclipse wide-field inverted microscope (Nikon 40 × 0.75 N.A. dry objective lens) equipped with Perfect Focus system and housed in a 37 °C chamber (Solent Scientific, UK) fed with 5% CO_2_. Multiple fields of view were selected at various XY coordinates, 3 Z slices were captured at a 1.25 μm spacing for HeLa cells and 1.8 μm spacing for HCT116 cells. Images were acquired using a Hamamatsu Orca Flash 4.0 camera (Hamamatsu Photonics, Japan), controlled by NIS-Elements software. For abscission time measurements, images were acquired every 10 minutes for 48 hours and abscission time was measured as the time from midbody formation to disappearance. For resolution timing for YFP-Lap2β positive bridges, images were acquired every 20 minutes for 72 hours, and resolution time was measured for time of appearance to disappearance of YFP-Lap2β positive intercellular bridges. For nuclear accumulation of GFP-NLS and analysis of anaphase defects, images were acquired every 5 minutes for 24 hours. Nuclear GFP signal was identified through co-localization with H2B-mCherry, and was normalized to the cytoplasmic signal. Signals were background corrected using measurements from adjacent regions. Measurements were taken 10 frames before and 50 frames after nuclear envelope breakdown, with cells depleted of CHMP7 serving as a positive control (56).

### Protein expression and purification

The human ALIX Bro1 domain (residues 1-359, Fig. 1*C*, *D*, Fig S1) and ALIX Bro1-V domains (residues 1-698, Fig. 1*B*) were expressed and purified as previously described with minor modifications (27, 57). A detailed description is provided by SI Materials and Methods. Plasmids for bacterial expression of ALIX proteins are available from the Addgene plasmid repository (www.addgene.org; see Table S2 for accession numbers).

### Crystallization and data collection

ALIX Bro1 crystallized in complex with CHMP4^A232^ or CHMP4C^T232^ peptides (residues 216-233; N-terminally acetylated) at 20°C from sitting drops that contained 1.2 μL of protein (200 μM ALIX Bro1 and 220 μM CHMP4C peptide) and 0.7 μL of reservoir solution (CHMP4C^A232^: 10% PEG 20,000, 100 mM MES, pH 6.5; CHMP4C^T232^: 15% PEG 8,000, 100 mM MES, pH 6.5, 200 mM sodium acetate). Crystals were flash frozen in nylon loops in cryo-protectant composed of reservoir solutions containing 30% glycerol. Data were collected remotely (58) (0.9794 Å wavelength, 100°K) at the Stanford Synchrotron Radiation Lightsource (SSRL) on beamline 12-2 using a Dectris Pilatus 6M detector. Data were integrated and scaled using AutoXDS (59-61). Both complexes crystallized in space group C121 with one ALIX-Bro1:CHMP4C complex in the asymmetric unit. The crystals diffracted to 1.91 Å (ALIX Bro1-CHMP4C^A232^) and 1.87 Å (ALIX Bro1-CHMP4C^T232^) resolution. For data collection and refinement statistics see Table S1.

### Structure determination and refinement

A model for molecular replacement was generated from the previously determined structure of the ALIX Bro1 -CHMP4C^A232^ complex (PDB 3C3R) (26) by removing the coordinates for the CHMP4C peptide and using the ALIX Bro1 structure as a search model (Phaser in PHENIX) (62, 63). CHMP4C helices were built *de novo* into the electron density for both structures using Coot (64), and further refined in Phenix (65, 66) using TLS refinement strategies (67). Final models had no outliers in the Ramachandran plot. Comparison of CHMP4C backbone atoms from both structures reveal that the N-termini (residues 221-229) superimpose with a 0.223 Å RMSD, whereas the C-termini (residues 229-233) superimpose with a 1.14 Å RMSD. Structures were analyzed and compared using PyMol (The PyMOL Molecular Graphics System, Version 1.3). Structure coordinates have been deposited in the RCSB Protein Databank with accession codes 5V3R and 5WA1 (ALIX Bro1-CHMP4C^A232^ and ALIX Bro1-CHMP4C^T232^, respectively).

### Cell growth assays

For assays examining the effect of low dose aphidicolin and reversine on cell growth cells were seeded at a density of 2.5 × 10^4^ cells per well of a 24 well plate in duplicate and a 12 well pate, treated 8 hours later with DMSO, 0.1 μM reversine or 30 nM aphidicolin, and left undisturbed for 2,4, (in 24 well plate) or 7 days (in 12 well plate), except for a media refresh after 4 days. Cell number was determined by manual counting at the indicated time points.

For analysis of cell growth following p53 depletion, cells were transduced with either control or p53 shRNA for 48 hours, then antibiotic selected for a further 48 hours. Transduced cells were seeded at a density of 2.5 × 10^4^ cells in a single well of a 24 well plate, cell number was determined at 2, 4, or 7 days. Cell number was determined by manual counting and cells were re-seeded in to 12 well plates on day 2, or 6 well plates on day 4.

### Karyotyping

HCT116^WT^ or HCT116^δCHMP4C^ cells were cultured for 48 hours in the presence of DMSO or 0.1 μM reversine and arrested in metaphase by overnight treatment with 50 ng/ml nocodazole. Cells were harvested, washed, treated in hypotonic buffer (10% FCS in ddH_2_O) for 30 minutes at 37°C and fixed in methanol:acetic acid (3:1 ratio). Fixation solution was replaced 4 times and spreads were produced by dropping 16 μL solution onto a glass slide from a height of 10 cm in a humid environment. Chromosomes from >90 metaphase-arrested cells were counted.

### Mitotic arrest assays

To examine the role of CHMP4C in mitotic spindle function HCT116^wT^ or HCT116^δCHMP4C^ cells were treated with 50ng/mL nocodazole, all cells were collected and analyzed by flow cytometry and western blotting. To examine the mitotic phosphorylation of CHMP4C, HeLa cells stably expressing indicated HA-CHMP4C constructs were treated with 2mM thymidine for 24 hours, washed, and treated with 50ng/mL Nocodazole overnight. All cells were collected and phosphorylation of HA-CHMP4C was determined by western blotting.

### Flow cytometry analysis

Cells treated overnight with 200 ng/mL mitomycin C, 50 ng/mL nocodazole or DMSO were harvested, washed in PBS and fixed in 1% paraformaldehyde. Cells were again washed and incubated with 50 μg/mL propidium iodide, 100 μg/mL RNase in PBS, and 0.1% Triton X-100. Cell cycle analysis profiles were acquired using FACS Canto II (BD Biosciences). 20,000 cells were counted per condition, and data were analyzed using FlowJo (Tree Start, Inc.). All gating was applied manually. For mitotic arrest experiments all conditions were performed in duplicate and samples were retained for immunoblotting analysis.

### Analysis of publicly available cancer genome-wide association studies in the UK Biobank

To determine if the CHMP4C rs35094336 variant is also associated with other cancer types we analyzed the publicly available data from 337,208 individuals in the UKBiobank engine. Results were obtained from Global Biobank Engine, Stanford, CA (URL: http://gbe.stanford.edu/) [October, 2017]. We used the following procedure to define cases and controls for cancer GWAS. Individual level ICD-10 codes from the UK Cancer Register, Data-Field 40006, and the National Health Service, Data-Field 41202 in the UK Biobank were mapped to the self-reported cancer codes, Data-Field 20001 as described previously (68). Positive associations are displayed in Table 1. A full review of the resource is available in reference (25).

## Acknowledgements

Funding for this project was provided by NIH grant R01 GM112080 (to WIS), Wellcome Trust grant WT102871MA (to JM-S), and support in conjunction with grant P30 CA042014 awarded to Huntsman Cancer Institute from the CRR Program at the Huntsman Cancer Institute (WIS, KSU). We thank the UK NIHR Comprehensive BRC at KCL for an equipment grant, and the Nikon Imaging Centre at KCL for technical support. DMW was supported in part by an American Cancer Society Postdoctoral Fellowship PF-14-102-01-CSM. Further support was provided by the Severo Ochoa Program grant SEV-2011-00067, a Sara Borrell Fellowship from the Instituto Carlos III and a Beatriu de Pinós fellowship from the Agency for Management of University and Research Grants (AGAUR) (to JMM), We acknowledge the Fluorescence Microscopy Core Facility at the University of Utah for access to imaging equipment, which was obtained using a NCRR Shared Equipment Grant 1S10RR024761-01. We thank Frank Whitby and Chris Hill for assistance with X-ray crystallography, and Scott Endicott at the University of Utah Peptide Synthesis core for peptide synthesis. Use of the Stanford Synchrotron Radiation Lightsource, SLAC National Accelerator Laboratory, is supported by the U.S. Department of Energy, Office of Science, Office of Basic Energy Sciences under Contract No. DE-AC02-76SF00515. The SSRL Structural Molecular Biology Program is supported by the DOE Office of Biological and Environmental Research, and by the National Institutes of Health, National Institute of General Medical Sciences (including P41GM103393). We thank Leandro Ventimiglia for sharing technical advice and cell lines. We also thank Jesus Gil (Imperial College London) and Pierre Vantourout (Kings College London) for the control and p53 shRNA and CRISPR Cas9 containing lentiviral packaging vector respectively. We thank Caroline Ogilvie (Kings College London) and Jennifer Pratt (Viapath), for guidance with the karyotyping experiments and the Manuel Rivas lab for making the UK Biobank resource available. The contents of this publication are solely the responsibility of the authors and do not necessarily represent the official views of NIGMS or NIH.

